# IPSE, a Parasite-Derived, Host Immunomodulatory Infiltrin Protein, Alleviates Resiniferatoxin-Induced Bladder Pain

**DOI:** 10.1101/2020.06.11.146829

**Authors:** Kenji Ishida, Evaristus C. Mbanefo, Loc Le, Olivia Lamanna, Luke F. Pennington, Julia C. Finkel, Theodore S. Jardetzky, Franco H. Falcone, Michael H. Hsieh

## Abstract

The transient receptor potential cation channel subfamily V member 1 (TRPV1) receptor is an important mediator of nociception and its expression is enriched in nociceptive neurons. TRPV1 signaling has been implicated in bladder pain and is a potential analgesic target. Resiniferatoxin is the most potent known agonist of TRPV1. Acute exposure of the rat bladder to resiniferatoxin has been demonstrated to result in pain-related freezing and licking behaviors that are alleviated by virally encoded IL-4. The interleukin-4-inducing principle of *Schistosoma mansoni* eggs (IPSE) is a powerful inducer of IL-4 secretion, and is also known to alter host cell transcription through a nuclear localization sequence-dependent mechanism. We previously reported that IPSE ameliorates ifosfamide-induced bladder pain in an IL-4- and nuclear localization sequence-dependent manner. We hypothesized that pre-administration of IPSE to resiniferatoxin-challenged mice would dampen pain-related behaviors. IPSE indeed lessened resiniferatoxin-triggered freezing behaviors in mice. This was a nuclear localization sequence-dependent phenomenon, since administration of a nuclear localization sequence mutant version of IPSE abrogated IPSE’s analgesic effect. In contrast, IPSE’s analgesic effect did not seem IL-4-dependent, since use of anti-IL-4 antibody in mice given both IPSE and resiniferatoxin did not dramatically affect freezing behaviors. RNA-Seq analysis of resiniferatoxin- and IPSE-exposed bladders revealed differential expression of TNF/NF-κb-related signaling pathway genes. *In vitro* testing of IPSE uptake by urothelial cells and TRPV1-expressing neuronal cells showed uptake by both cell types. Thus, IPSE’s nuclear localization sequence-dependent therapeutic effects on TRPV1-mediated bladder pain may act on TRPV1-expressing neurons and/or may rely upon urothelial mechanisms.

## Introduction

The bladder is a heavily innervated organ [1]. The high density of afferent nerve endings in the bladder partly accounts for its sensitivity to noxious stimuli (reviewed by de Groat and Yoshimura [2]). Diverse stimuli cause bladder-based nociception, including urinary tract infections, catheterization, surgical manipulation, ureteral stents, hemorrhagic cystitis, and bladder pain syndromes such as interstitial cystitis [2]. Despite the importance of bladder-based nociception in clinical medicine, there are few therapeutic options that directly target afferent nerve endings in the bladder.

One potential set of therapeutics for bladder pain are the proteins known as the Interleukin-4-inducing principle of *Schistosoma mansoni* eggs (IPSE) [3]. As the name indicates, IPSE is a potent inducer of IL-4 secretion by host cells. IPSE also features a nuclear localization sequence (NLS) which facilitates its entry into host cell nuclei and subsequent modulation of transcription [4,5]. Macedo et al. have reported that administration of IL-4 to mice with ifosfamide-induced hemorrhagic cystitis alleviates bladder injury [6]. This led us to test IPSE in this model. A single dose of IPSE reduced spontaneous pain behaviors in ifosfamide-challenged mice in an IL-4- and NLS-dependent manner [7].

Other investigators have reported that administration of virally-encoded IL-4 reduces resiniferatoxin-induced, bladder pain-related behaviors [8]. This led us to hypothesize that IPSE may likewise dampen bladder pain caused by resiniferatoxin. Herein we describe the ability of IPSE to ameliorate resiniferatoxin-triggered bladder pain behaviors. This property of IPSE is NLS-dependent, and possibly weakly IL-4-dependent. RNA-Seq analysis of resiniferatoxin- and IPSE-exposed bladders indicates that IPSE reduces gene expression related to the TNF signaling via NF-κB pathway. These effects occur in the context of uptake of IPSE by both urothelial and neuronal cells.

## Materials and Methods

### Study approval

All animal work was conducted according to relevant U.S. and international guidelines. Specifically, animal experimental work was reviewed and approved as protocol 14-03 by the Institutional Animal Care and Use Committee of the Biomedical Research Institute (Rockville, Maryland, USA). Our Institutional Animal Care and Use Committee guidelines comply with the U.S. Public Health Service Policy on Humane Care and Use of Laboratory Animals.

### Mice

Female 6- to 8-wk-old C57BL/6 mice (Charles River Laboratories, Wilmington, MA, USA) were housed under 12 h light-dark cycles in temperature-controlled holding rooms with unlimited access to dry mouse chow and water. Newly received mice were acclimated to the animal facility for at least one week prior to experimental use.

### IPSE protein production and labeling

Recombinant H06 H-IPSE (one of the major *Schistosoma haematobium* IPSE orthologs) and an NLS mutant of H06 H-IPSE were produced in HEK293-6E cells as previously described [9]. H06 IPSE was conjugated to Alexa Fluor 488 using a Alexa Fluor 488 antibody labeling kit (Thermofisher Scientific, Waltham, MA) according to manufacturer instructions; however, the pH was kept at 7.4 throughout the reaction to enrich for labeling of the terminal amine (pKA of 7.4). The efficiency of conjugation was confirmed by Nanodrop. The typical degree of labeling was one mole of dye per mole of IPSE, which suggested IPSE was only labeled on the terminal amine. Low labeling efficiently minimized the potential interference of the dye with IPSE’s functional domains.

### IPSE administration

One day prior to resiniferatoxin or vehicle challenge, mice underwent tail vein injection with phosphate-buffered saline or 25 μg of H06 H-IPSE (or its NLS mutant) in phosphate-buffered saline.

### Recombinant IL-4 administration

Recombinant mouse IL-4 was obtained from Peprotech Laboratories (Rocky Hill, NJ, USA). A subset of mice underwent i.p. injection with 10 ng of IL-4 one hour prior to resiniferatoxin challenge.

### Anti-IL-4 antibody administration

Neutralizing anti-IL-4 antibody (11B11 clone) was purchased from BioXcell (West Lebanon, NH, USA). A subset of mice underwent i.p. injection with 100 μg of anti-IL-4 antibody 30 minutes before resiniferatoxin challenge.

### Resiniferatoxin administration and assessment of freezing behavior

Mice were anesthetized, treated, and evaluated one by one. Anesthesia was achieved using vaporized isoflurane and mice kept on a heating blanket to maintain body temperature. Lubricated sterile catheters (Excel Safelet Cath 24G x 3/4“) attached to 1 mL-syringes were gently inserted into the mouse urethra. Phosphate-buffered saline (50 μL), vehicle (50 μL of 10% v/v ethanol, 10% v/v Tween 80 in 80% v/v phosphate-buffered saline) or resiniferatoxin (3 μM in 50 μL of 10% v/v ethanol, 10% v/v Tween 80 in 80% v/v phosphate-buffered saline) was slowly pushed into the mouse bladder and held in place for 1 minute.

Mice were withdrawn from the anesthesia and allowed to recover from the effect of anesthesia by leaving them on the warm pad for 5 min. Subsequently, mice were transferred to a transparent cage and a continuous video footage were recorded for 15 minutes following resiniferatoxin administration. Freezing behaviors were scored for each individual mouse over 5 minute periods (5 min, 10 min and 15 min) in a blinded fashion [8].

### RNA purification

RNA was isolated from mouse bladders using TRIzol Reagent and PureLink RNA Mini Kit (Invitrogen), according to manufacturers’ instructions. Briefly, aseptically excised bladders were homogenized in 1 ml TRIzol Reagent by bead-beating using ceramic beads (Omni International) and a mini-bead beater (Biospec). Following a 5-min incubation, 0.2 ml chloroform was added and again incubated for 3 min before centrifugation at 12,000 × g for 15 min to separate homogenates into aqueous and organic phases. The aqueous supernatant (∼400ul) was mixed with an equal volume of 70% ethanol before binding the mixture to RNA binding columns by centrifugation. On-column DNase digestion (Invitrogen) was performed for 30 minutes, following the manufacturer’s protocols. After column washes and drying, RNA was eluted in RNase-free water, quantified and its quality checked using a NanoDrop 1000 spectrophotometer (Thermo Scientific) and Bioanalyzer 2100 (Agilent).

### RNA sequencing and RNA-seq analysis pipeline

RNA sequencing was performed using the Illumina-HiSeq 4000 NGS platform at a depth of >20 million reads. Analyses were conducted using the RNA analysis tools of the Galaxy platform (https://usegalaxy.org). Raw sequence reads were aligned to the mouse genome (mm10) by HISAT2 (version 2.1.0+galaxy4). The resulting alignment files, along with the most recent mouse genome annotation file in the Illumina iGenomes UCSC mm10 mouse genome collection (http://igenomes.illumina.com.s3-website-us-east-1.amazonaws.com/Mus_musculus/UCSC/mm10/Mus_musculus_UCSC_mm10.tar.gz), were used as the input for HTSeq-count (version 0.9.1). DESeq2 (Galaxy version 2.11.40.6+galaxy1; DESeq2 version 1.22.1) was used to determine differentially expressed genes across all treatment groups. PCA plots were also generated by DESeq2.

### Functional and pathway analysis, statistics and plots

Treatment-to-pathway association was performed with the Gene Set Enrichment Analysis (GSEA) software (version 4.0.3)(https://www.gsea-msigdb.org/gsea/index.jsp), using the DESeq2 normalized read counts file from which genes that showed zero read counts for any sample were removed, hallmark gene set (version 7.1)(ftp://ftp.broadinstitute.org/distribution/gsea/gene_sets/h.all.v7.1.symbols.gmt), mouse gene symbol remapping file (version 7.1)(ftp://ftp.broadinstitute.org/distribution/gsea/annotations_versioned/Mouse_Gene_Symbol_Remapping_to_Human_Orthologs_MSigDB.v7.1.chip), with “Permutation type” set to “Gene_set”, “Create GCT files” set to “True”, and other analysis options set to default values [10]. The volcano plot was generated using the EnhancedVolcano software package (version 1.4.0 from the bioconda distribution channel) for R [11]. The heat maps were generated using the Morpheus software (https://software.broadinstitute.org/morpheus). Other data analyses and plots were generated using GraphPad Prism v 6.00, and ggplot2 and plotly packages in R. For comparisons among groups, one-way analysis of variance (ANOVA) was performed and if significant, was followed by post hoc Student t-tests for pairwise comparisons after confirming a normal distribution. Plotted data show individual data points with error bars representing means and standard deviation.

### Endocytosis assays

Cath.a mouse brain-derived neuronal cells (ATCC CRL-11179) were obtained from ATCC (Manassas, VA) and were grown in RPMI-1640 (Sigma-Aldrich, St. Louis, MO) with 8% horse serum (Sigma-Aldrich, St. Louis, MO) and 4% fetal bovine serum (Sigma-Aldrich, St. Louis, MO). HCV-29 human derived urothelial cells were obtained as a gift from Paul Brindley and grown in MEM (Thermofisher Scientific, Waltham, MA) with 10% fetal bovine serum (Sigma-Aldrich, St. Louis, MO). For internalization assays, floating cells and adherent cells (released via 0.12% trypsin (Sigma-Aldrich, St. Louis, MO) without EDTA) were washed in fresh medium, and aliquoted into 24 well plates at 200,000 cells/mL in 1 mL. The cells were incubated with Alexa 488-labeled H06 at 1 μg/mL or Alexa 488-labeled transferrin at 4 μg/mL (Thermofisher Scientific, Waltham, MA) for 16 hours at 37° C. Cells were released via 0.12% trypsin without EDTA and washed 3 times with PBS (Sigma-Aldrich, St. Louis, MO). 0.4% trypan blue (Thermofisher Scientific, Waltham, MA) was added to the cells (1:4) to quench extracellular Alexa 488 signal. The cells were analyzed by flow cytometry (Beckman Coulter, CytoFLEX) to isolate the intracellular Alexa 488 signal. Data was analyzed using FlowJo and GraphPad.

## Results

### IPSE reduces resiniferatoxin-induced, bladder pain-associated behaviors in an IL-4- and nuclear localization sequence-dependent fashion

When mice were given intravesical resiniferatoxin, they exhibited a significant increase in pain-associated freezing behaviors (Figure 1A). Administration of a single intravenous dose of IPSE 24 hours prior to resiniferatoxin challenge resulted in significantly decreased freezing episodes. However, IPSE did not bring freezing episodes down to the levels of vehicle-treated mice (e.g., no resiniferatoxin exposure).

**Figure 1.**
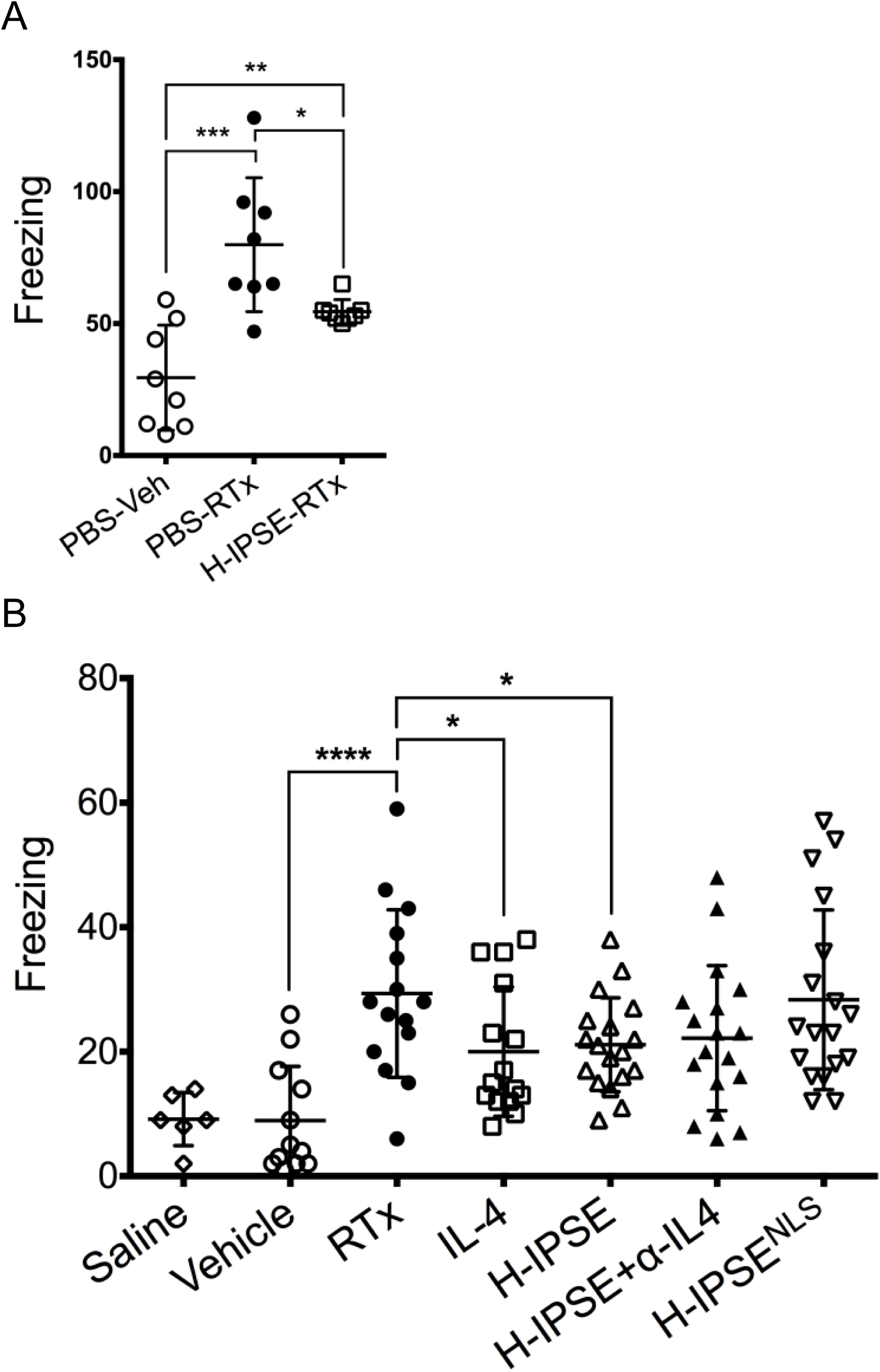
IPSE reduces resiniferatoxin-induced, pain-related freezing behaviors in an IL-4- and nuclear localization sequence-dependent manner. A, mice were administered intravenous phosphate-buffered saline (PBS) followed by intravesical PBS/Tween/ethanol vehicle (“PBS-Veh”), intravenous PBS with intravesical resiniferatoxin in vehicle (“PBS-RTx”), or one intravenous dose of the H06 H-IPSE ortholog of IPSE 24 hours before intravesical resiniferatoxin in vehicle (“H-IPSE-RTx”). B. mice were given intravenous PBS and intravesical PBS (“saline”), intravenous PBS and intravesical PBS/Tween/ethanol vehicle (“Vehicle”), intravenous PBS and intravesical resiniferatoxin in vehicle (“RTx”), recombinant IL-4 given intraperitoneally followed by intravesical resiniferatoxin in vehicle (“IL-4”), the H06 H-IPSE ortholog of IPSE given intravenously 24 hours before intravesical PBS with resiniferatoxin in vehicle (“H-IPSE”), the H06 H-IPSE ortholog of IPSE 24 hours given intravenously and anti-IL-4 antibody given intraperitoneally 30 minutes before resiniferatoxin in vehicle administered intravesically (“H-IPSE+ α-IL4”), or a nuclear localization sequence (NLS) mutant of H06 H-IPSE given intravenously 24 hours before resiniferatoxin in vehicle administered intravesically (“H-IPSE^NLS^”).

In an independent set of experiments we then tested the ability of an NLS mutant of IPSE, as well as IPSE combined with anti-IL-4 antibody, to reduce resiniferatoxin-induced pain behaviors compared to wild type IPSE and recombinant IL-4 (positive control) (Figure 1B). H06-IPSE had similar effects to recombinant IL-4. The treatment with an IL-4 blocking antibody half an hour before resiniferatoxin administration and 24 hours after IPSE treatment did not appear to have a strong effect on IPSE’s lessening of resiniferatoxin-induced, pain-related freezing behaviors, suggesting that the effects of IPSE are not dependent on IL-4 release. In contrast to wild type IPSE, the NLS mutant of IPSE did not seem to exert an analgesic effect on resiniferatoxin-exposed mice.

### IPSE decreases resiniferatoxin-induced bladder expression of genes associated with TNF signaling via NF-κb

We next sought to determine the effects of IPSE and resiniferatoxin treatment on bladder transcription. Mice were administered H06 H-IPSE and resiniferatoxin, and their bladders harvested for RNA-Seq analysis. Principal component analysis (PCA) confirmed that resiniferatoxin-treated bladders clustered distinctly from vehicle-treated bladders (Figure 2). Likewise, PCA of IPSE combined with resiniferatoxin versus resiniferatoxin only-treated bladders also showed distinct clustering patterns, albeit less so (Figure 2). By analysis with DESeq2, we found 219 differentially expressed genes (adjusted p-value < 0.1), 592 genes with an absolute value of Log2 fold change > 0.322 (greater than 1.25-fold change in either direction), and 129 genes satisfying both conditions in the comparison between IPSE combined with resiniferatoxin versus resiniferatoxin treatment groups (Figure 3; Supplementary Table 1). To determine whether IPSE could restore or rescue the expression of genes perturbed by the resiniferatoxin treatment, we performed the following pairwise comparisons: resiniferatoxin versus vehicle (RTXvsVeh; Supplementary Table 2), IPSE versus resiniferatoxin (H06vsRTX; Supplementary Table 1), and IPSE versus vehicle (H06vsVeh; Supplementary Table 3). In the first scenario, we filtered for genes whose expression was increased in the resiniferatoxin treatment compared to both vehicle and H06 treatment using the following conditions: Log2 fold change > 0.322 with adjusted p-value < 0.1 in RTXvsVeh; Log2 fold change < −0.322 with adjusted p-value < 0.1 in H06vsRTX; and adjusted p-value >= 0.1 in H06vsVeh. The filtering for this scenario yielded 21 genes (Supplementary Table 4). Similarly, in the second scenario, we filtered for genes whose expression was decreased in the resiniferatoxin treatment compared to both vehicle and H06 treatment using the following conditions: Log2 fold change < −0.322 with adjusted p-value < 0.1 in RTXvsVeh; Log2 fold change > 0.322 with adjusted p-value < 0.1 in H06vsRTX; and adjusted p-value >= 0.1 in H06vsVeh. This scenario yielded 23 genes (Supplementary Table 5).

**Figure 2.**
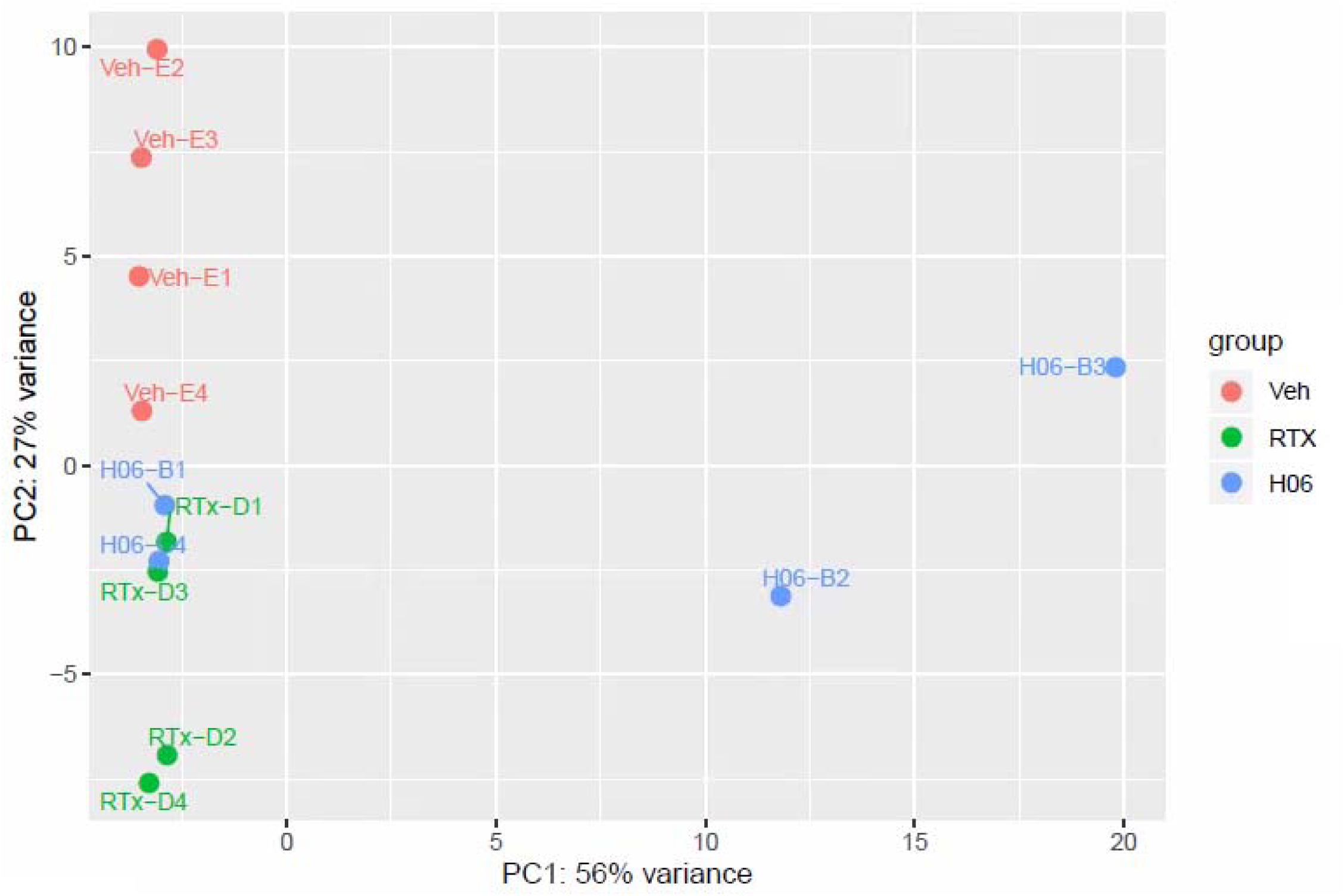
Principal component analysis of resiniferatoxin- and IPSE-treated bladder gene expression. Principal component analysis showed homogeneous clustering of gene expression among resiniferatoxin-treated mice (green symbols labeled with “RTx-D”) and vehicle-treated mice (red symbols labeled with “Veh-E”). There was some overlap of gene expression among resiniferatoxin-treated mice and mice treated with both H06 H-IPSE and resiniferatoxin (blue symbols labeled with “H06-B”).

**Figure 3.**
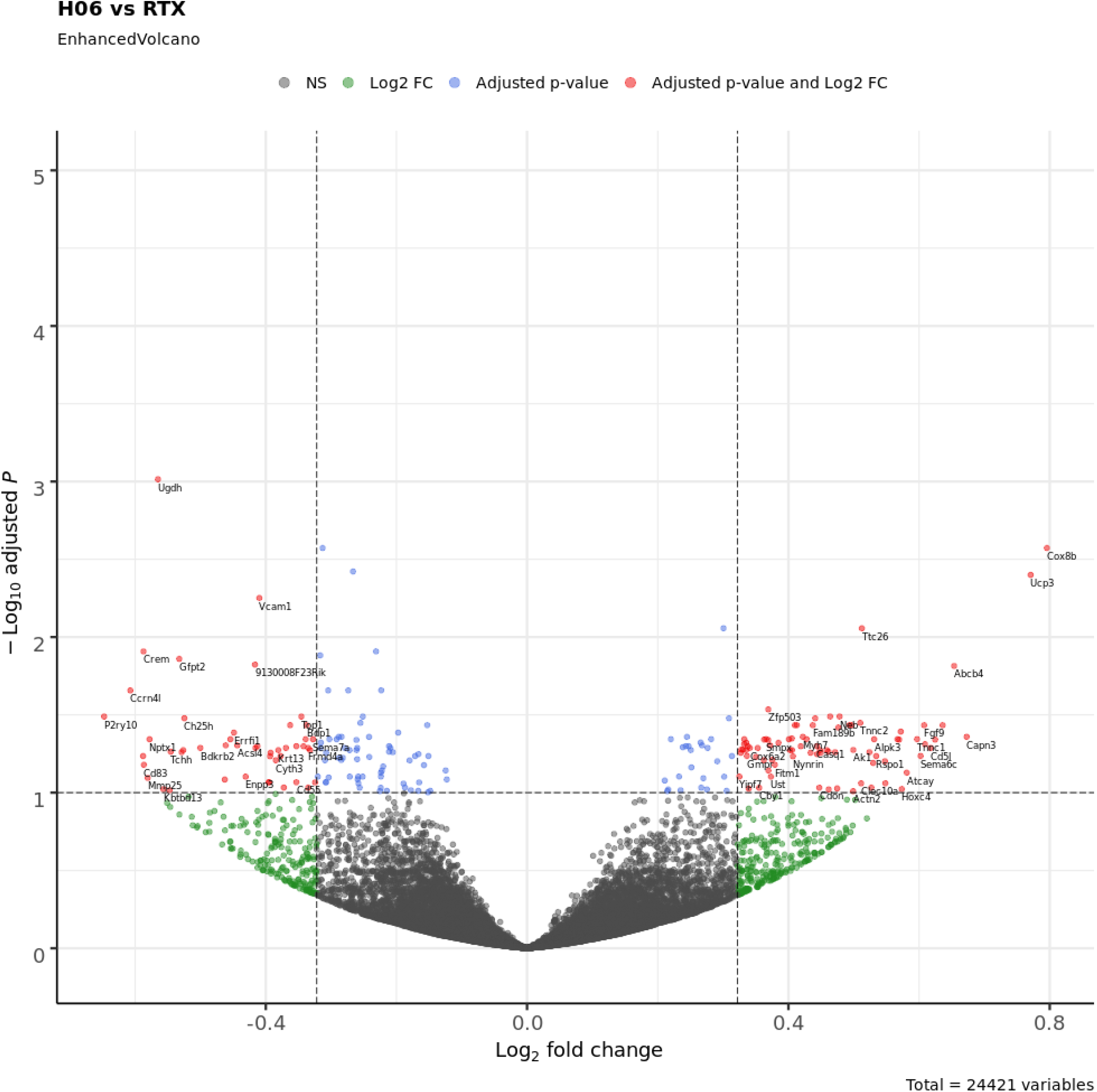
Volcano plot showing differentially expressed genes between bladders treated with IPSE and resiniferatoxin (H06) versus bladders treated with resiniferatoxin alone (RTX). The cutoff value for the adjusted p-value was set at < 0.1, and the cutoff for the absolute value of the Log2 fold change was set at > 0.322 (1.25-fold in either direction). Blue dots represent genes satisfying the adjusted p-value cutoff. Green dots represent genes satisfying the Log2 fold change cutoff. Red dots represent genes satisfying both of these cutoff conditions and are labeled with the corresponding gene symbols. Gray dots represent genes that do not satisfy either condition (NS, not significant).

Using the normalized read counts file from DESeq2 (Supplementary Table 6) processed to remove genes for which any sample showed a zero read count, Gene Set Enrichment Analysis (GSEA) software, and Morpheus software, we then generated a heat map of differential gene expression in bladders treated with H06 H-IPSE and resiniferatoxin versus resiniferatoxin alone (Figure 4; Supplementary Table 7). Among the 50 hallmark gene sets in the Molecular Signatures Database (https://www.gsea-msigdb.org/gsea/msigdb/collections.jsp), many gene sets with a false discovery rate (FDR) q-value < 0.05 were enriched in the resiniferatoxin-alone treatment, including TNF signaling via NF-κB, inflammatory response, allograft rejection, interferon gamma response, and IL6/JAK-STAT3 signaling (Supplementary Table 8). Of these, the TNF signaling via NF-κB gene set showed the greatest normalized enrichment score in terms of absolute magnitude. The enrichment plot for the TNF signaling via NF-κB gene set shows a negative peak in the enrichment score, indicating a greater correlation of this gene set to the resiniferatoxin-alone treatment when compared to the IPSE combined with resiniferatoxin treatment (Figure 5). Notably, IL6 and IL1B were more strongly associated with the resiniferatoxin-alone treatment compared to the IPSE-resiniferatoxin treatment (Figure 6; Supplemental Table 9).

**Figure 4.**
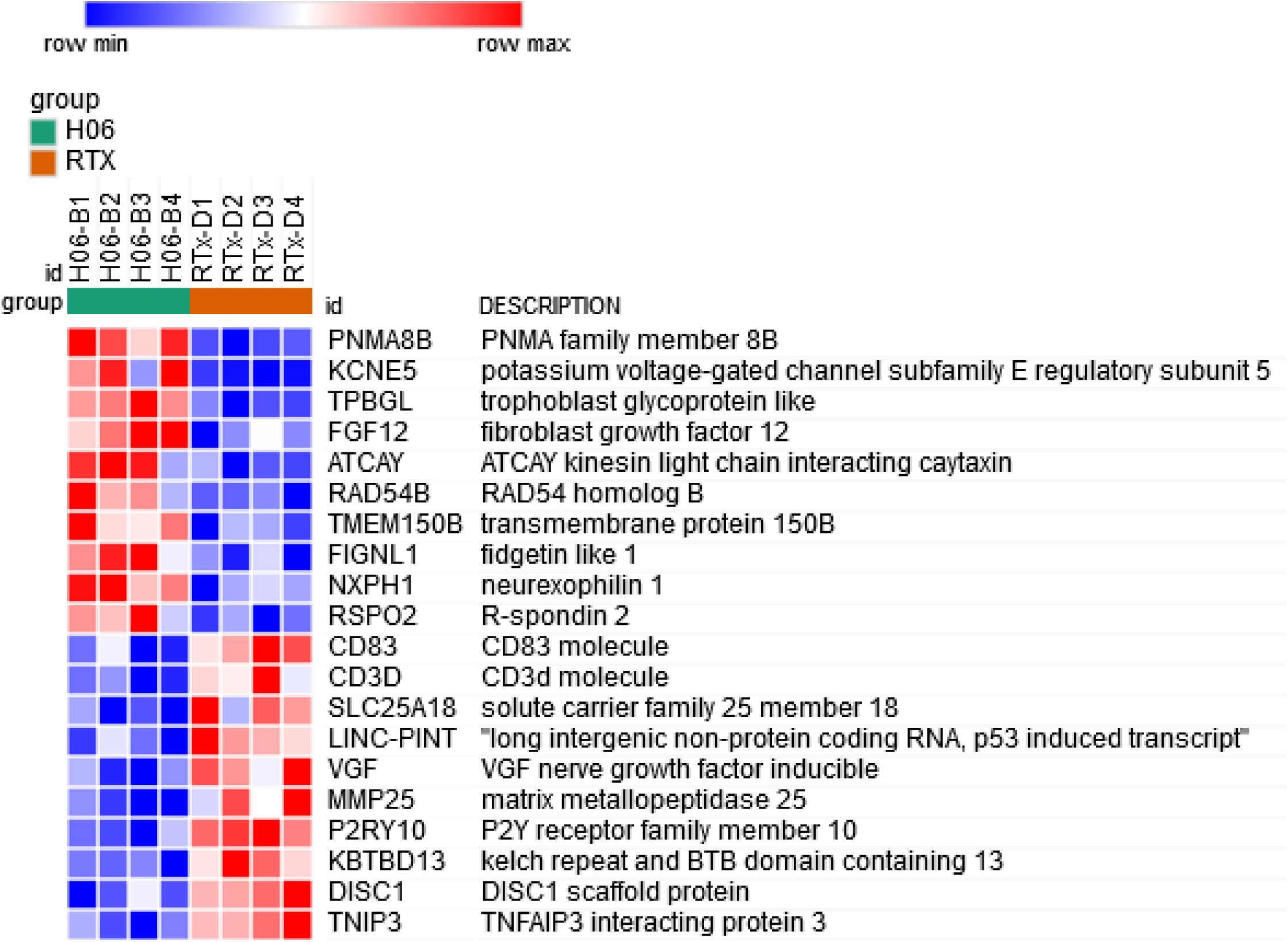
Heat map of the top and bottom 10 genes from the 50 hallmark gene sets from the Molecular Signatures Database with enriched differential expression in bladders exposed to IPSE combined with resiniferatoxin or resiniferatoxin alone. Each column shows gene expression for an individual mouse bladder. Green and brown column coloring indicates IPSE combined with resiniferatoxin (H06) versus resiniferatoxin only (RTX)-treated bladders, respectively. Genes are sorted by signal-to-noise scores, and their symbols and names are listed in rows. Darkest blue to darkest red coloring represents lowest to highest gene expression, respectively, based on normalized read counts.

**Figure 5.**
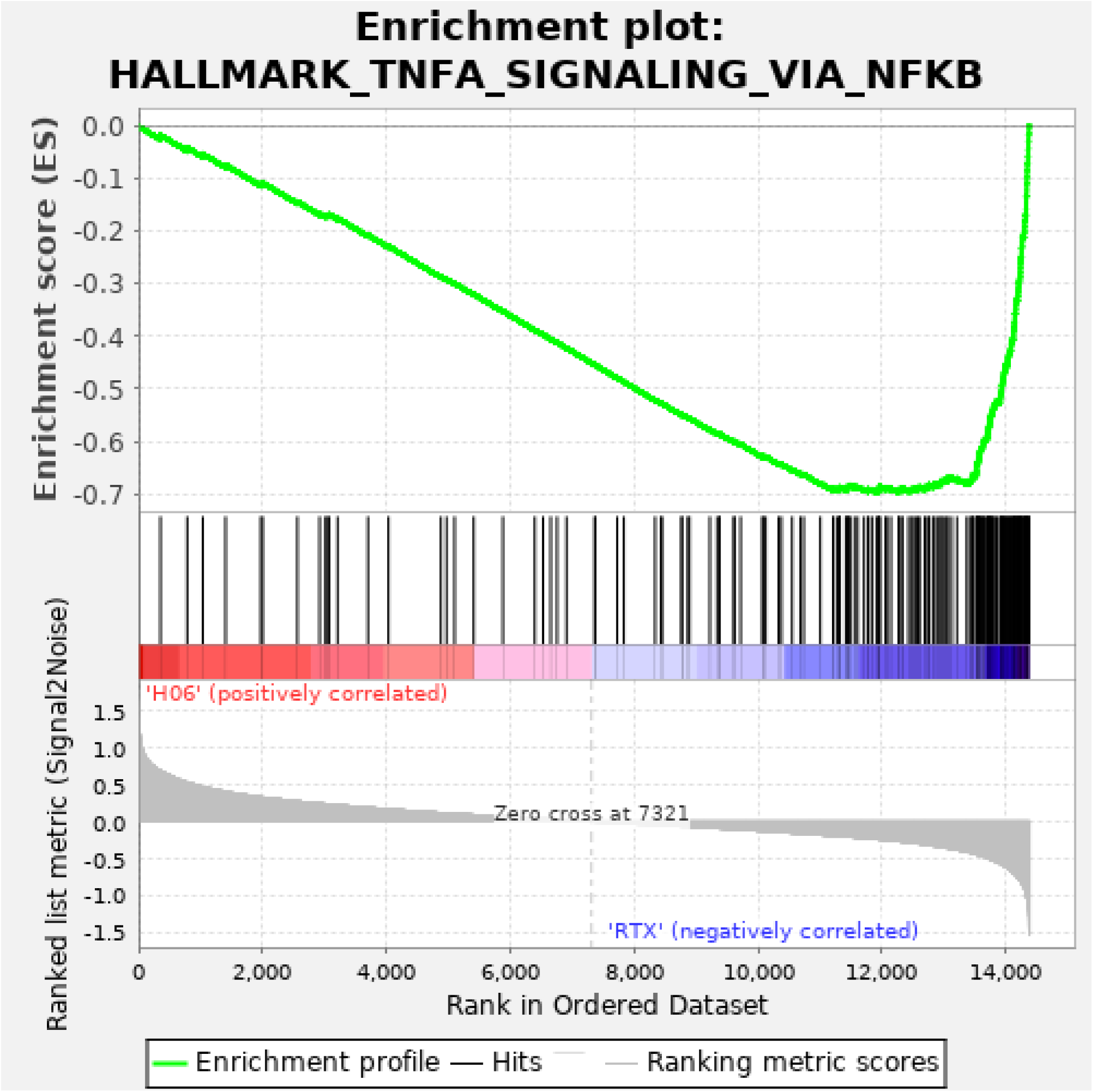
Enrichment plot for the TNF signaling via NF-κB pathway. Top panel, green line indicates running enrichment score for the TNF signaling via NF-κB pathway as the gene set enrichment analysis walks down the ranked list of genes. Middle panel depicts where the members of the TNF signaling via NF-κB pathway gene set appear in the ranked list of genes. Bottom panel shows the value of the ranking metric moving down the list of ranked genes. Positive values indicate correlation with the first phenotype (H06; H-IPSE combined with resiniferatoxin) and negative values indicate correlation with the second phenotype (RTX; resiniferatoxin alone).

**Figure 6.**
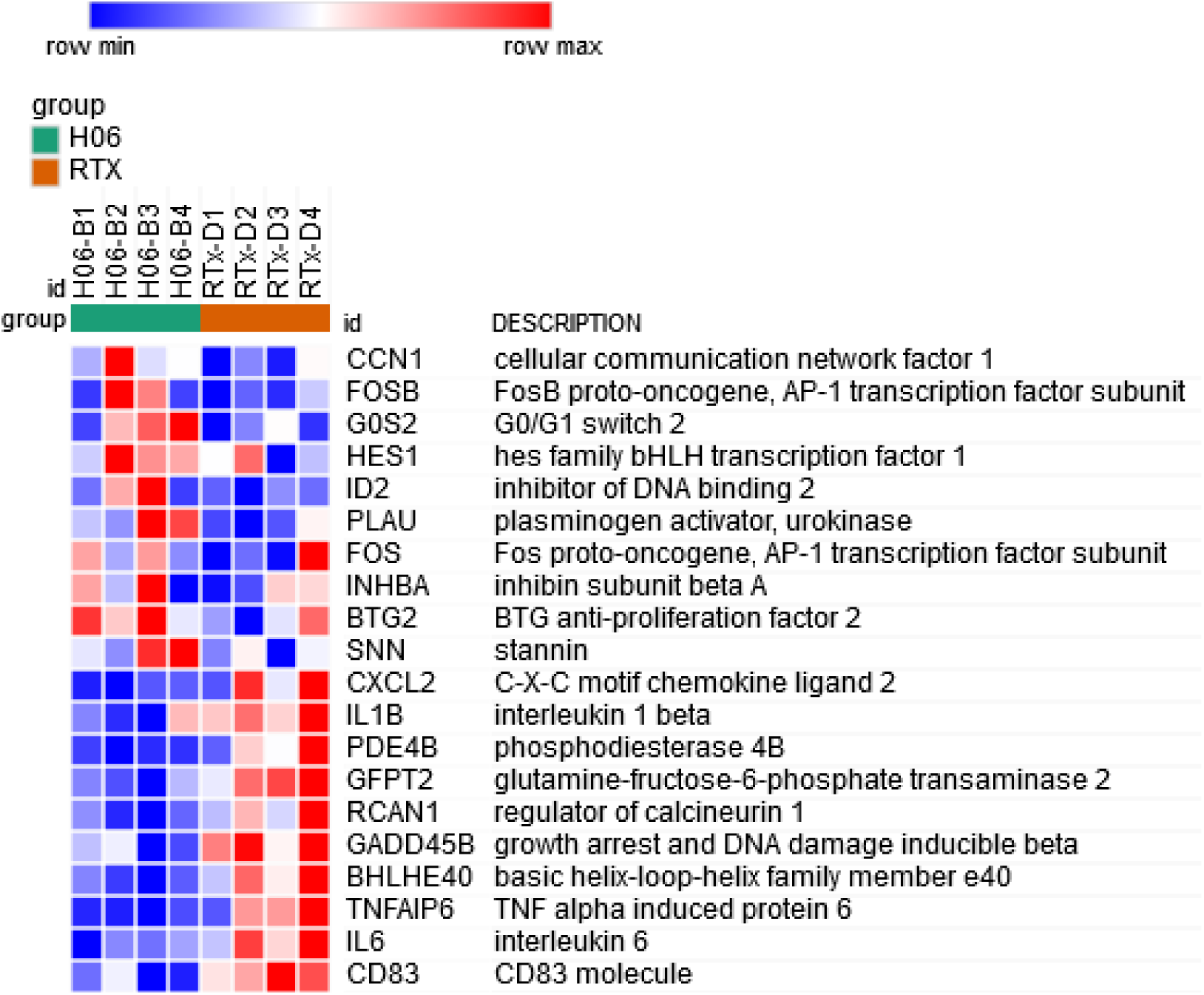
Heat map of the top and bottom 10 gene members of the TNF signaling via NF-κB pathway with enriched differential expression in bladders exposed to IPSE combined with resiniferatoxin or resiniferatoxin alone. Each column shows gene expression for an individual mouse bladder. Green and brown column coloring indicates IPSE combined with resiniferatoxin (H06) versus resiniferatoxin only (RTX)-treated bladders, respectively. Genes are sorted by signal-to-noise scores, and their symbols and names are listed in rows. Darkest blue to darkest red coloring represents lowest to highest gene expression, respectively, based on normalized read counts.

### IPSE is taken up by both neuronal and urothelial cells via endocytosis

Resiniferatoxin is the most potent known agonist of the transient receptor potential cation channel subfamily V member 1 (TRPV1) receptor. Given that expression of this receptor is enriched in afferent neurons, we hypothesized that IPSE may mediate some or most of its effects on resiniferatoxin-induced pain through neuronal endocytosis and downstream modulation of neuronal transcription. To test this hypothesis, we sought to measure endocytosis of IPSE by Cath.a mouse neuronal cells versus HCV-29 urothelial cells (Figure 7). Despite differences in endocytosis of transferrin control between Cath.a cells and HCV-29 cells, we found that Cath.a endocytosis of IPSE was similar to that of urothelial cells, which are known to take up IPSE [12]. This lends credence to the theory that IPSE may exert some of its analgesic effects through neuronal mechanisms, but also supports a possible urothelial role in IPSE’s bladder analgesic properties.

**Figure 7.**
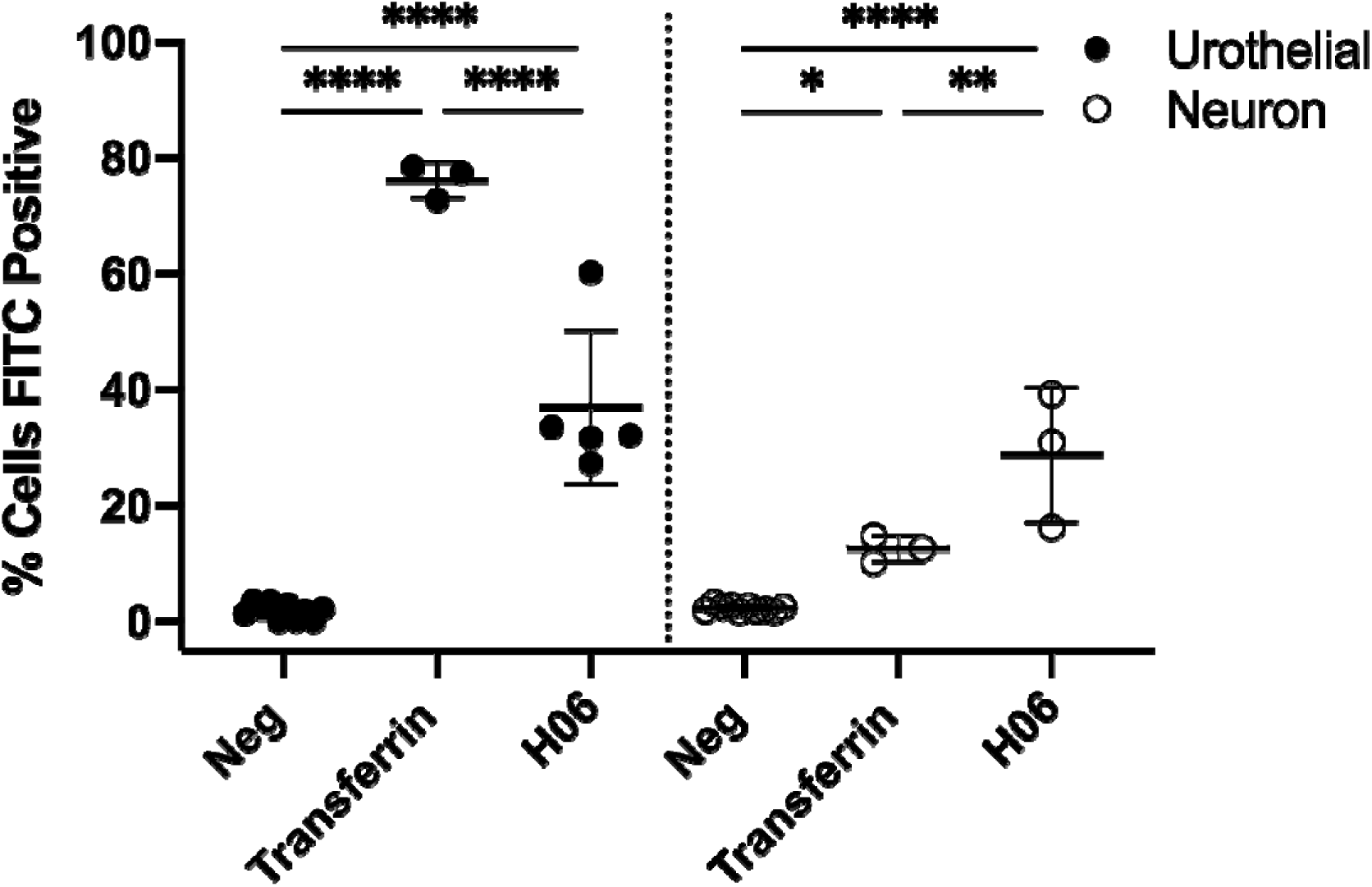
Internalization of IPSE by neuronal and urothelial cells. Cath.a mouse neuronal and HCV-29 human urothelial cells were incubated for 16 hrs with Alexa 488 conjugated H06 H-IPSE (1 μg/ml) or transferrin (4 μg/ml) and analyzed by flow cytometry after trypan blue extracellular signal quenching. Data is representative of 2 experiments. *p=0.0139, **p=0.0025, ****p<0.0001.

## Discussion

Bladder pain can be caused by infection, inflammation, instrumentation, or poorly understood conditions such as bladder pain syndrome. Regardless of etiology, there is a lack of therapeutics that target bladder pain. One study estimates that 3.3-7.9 million women in the US suffer from bladder pain symptoms [13]. In the United States alone, interstitial cystitis/bladder pain syndrome costs ∼$20–40 billion per annum to treat [14]. The high costs of this condition reflect limitations in available efficacious treatments. Although the pathophysiology of interstitial cystitis/bladder pain syndrome is not completely understood, prior bladder infection, stress, and changes to neural pathways may play roles in the associated pain [2].

Novel analgesics have attempted to target TRPV1 (capsaicin receptor)-expressing afferent neurons. Interestingly, TRPV1-expressing neurons have been implicated in chemotherapy-induced hemorrhagic cystitis as well as inflammation associated with interstitial cystitis/bladder pain syndrome [15,16]. Hence, analgesics targeting TRPV1-expressing neurons may be a promising therapeutic approach for bladder pain caused by disparate noxious stimuli.

Parasite-derived molecules hold promise as non-opioid analgesics. Parasites have closely co-evolved with humans, and in the process have evolved the ability to produce molecules which modulate host inflammation to prevent parasite death. This observation has led to “helminth therapy”, including administration of helminth eggs to patients with inflammatory bowel disease to decrease disease flares and symptoms [17]. A likely safer approach to helminth therapy would be to generate recombinant parasite-derived proteins and administer these single proteins to patients based on known disease mechanisms.

One set of parasite proteins with significant therapeutic potential is the group of homologs of the Interleukin-4 inducing Principle of Schistosoma mansoni Eggs (IPSE) [18]. IPSE, also known as α-1 [3] and has multiple host immune modulatory functions. Firstly, IPSE ligates Fcε receptor-bound IgE on the surface of basophils and mast cells to induce IL-4 secretion [7,19,20]. It is also able to bind to immunoglobulins on the surface of B regulatory cells (Bregs) and thereby activate these cells [21]. The *S. mansoni* ortholog of IPSE called *S. mansoni* chemokine-binding protein (smCKBP) can neutralize chemokines [22]. Finally, IPSE contains a nuclear localization sequence which directs the protein to host cell nuclei [4,9], where it modulates transcription [5,12].

IPSE’s IL-4-influencing properties led us to test its ability to lessen IL-4-dependent, ifosfamide-induced hemorrhagic cystitis [5,7,23]. Besides verifying that IPSE indeed could dampen ifosfamide-triggered hemorrhagic cystitis in an IL-4-dependent manner, we also found that many of IPSE’s effects in this model relied upon an intact nuclear localization sequence [7,23]. In a subsequent RNA-Seq-based analysis, we confirmed that gene transcription related to TNF signaling is upregulated in ifosfamide-induced hemorrhagic cystitis [24], as reported by others using alternative experimental approaches [25,26]. Moreover, through this analysis we discovered that IPSE reduces expression of ifosfamide-induced genes related to the TNF pathway.

TNF signaling has also been implicated in promotion of resiniferatoxin-induced nociception [27]. Resiniferatoxin is the most potent known agonist for the nociception-associated TRPV1 receptor. TRPV1 stimulation by resiniferatoxin causes this ion channel to become permeable to cations, including calcium. The influx of calcium and other cations causes TRPV1-expressing neurons to depolarize, transmitting strong nociceptive signals. This acute stimulation is followed by desensitization and analgesia, in part because nerve endings die from calcium overload [28,29].

Oguchi et al. reported that resiniferatoxin-induced bladder pain could be alleviated by virally delivered IL-4 [8]. Considering IPSE’s IL-4-inducing properties, we postulated that IPSE could also lessen resiniferatoxin-triggered bladder pain through IL-4-related pathways. Although we did not definitively confirm IPSE could decrease resiniferatoxin-induced bladder pain via IL-4-dependent signaling, we did verify IPSE exerts analgesia through nuclear localization sequence-dependent mechanisms (Figure 1B), similar to our observations in the ifosfamide-induced hemorrhagic cystitis model [7,23]. Furthermore, bladder transcriptional profiling revealed a role for TNF pathways in resiniferatoxin-triggered bladder pain (Figures 4 and 5 and Table 1), parallel to our findings in ifosfamide-induced hemorrhagic cystitis [5]. Lastly, we discovered that IPSE decreases gene transcription of TNF-associated pathways induced by resiniferatoxin (Figures 4 and 5 and Table 1), again mirroring our observations using the ifosfamide-triggered model of hemorrhagic cystitis-associated bladder pain [5].

Our work has noteworthy limitations. Although a single dose of IPSE prior to resiniferatoxin exposure greatly decreased bladder pain-associated behaviors, it did not abolish them completely (Figure 1). In addition, IPSE did not alleviate licking, another set of resiniferatoxin-induced nociceptive behaviors (data not shown). Future work will examine the effects of repeated doses of H06 H-IPSE, as well as other wild type and mutant orthologs of IPSE. Despite an apparent analgesic phenotype, H06 H-IPSE did not lead to radical changes in the transcriptome of the resiniferatoxin-exposed bladder (Figure 2). However, the observed differential expression of TNF-associated genes is consistent with known effects of resiniferatoxin (Figures 4 and 5 and Table 1), and are also well-aligned with our observations in ifosfamide-induced hemorrhagic cystitis [5] [27]. Finally, it is possible that trypsinization of the neuronal and urothelial cells prior to IPSE uptake experiments may affect cellular endocytosis. Nonetheless, HCV-29 urothelial cells demonstrated 80% transferrin uptake 16 hours following trypsinization, suggesting that transferrin receptor function recovers well after trypsin exposure. Cath.a neuronal cells primarily grow buoyant in suspension, and only a minority of cells are adherent and require trypsinization to release them. Assuming the majority of buoyant Cath.a cells have intact transferrin receptors (due to lack of exposure to trypsin), the low transferrin and higher IPSE uptake indicates that Cath.a cell endocytosis of IPSE is not clearly adversely affected by trypsin exposure.

In summary, a single intravenous dose of H06 H-IPSE ameliorates bladder pain induced by resiniferatoxin, the most potent known agonist for TRPV1, an ion channel widely expressed by nociceptive neurons. H06 H-IPSE exerts this effect through nuclear localization sequence-linked pathways and does so in the context of endocytosis by both neurons and urothelial cells. This indicates that H06 H-IPSE’s analgesic features may depend on the molecule’s multiple host modulatory functions. Additionally, these functions may act upon neurons, but may also be executed through effects on other cell types that express TRPV1 or that modulate neuronal properties. For example, human leukocytes have been reported to express TRPV1 [30], as well as urothelial cells (reviewed by Andersson [31]). Ongoing efforts will help identify IPSE’s mechanisms of effect on TRPV1-associated nociception and may contribute to development of IPSE as a novel analgesic.

## Acknowledgement

We thank Paul Brindley for the gift of HCV-29 cells. We are grateful for support of this work by NIH R01DK113504.

## Author Contributions

KI: performed experiments, conducted data analysis, assisted with manuscript editing

ECM: designed and performed experiments, conducted data analysis, assisted with manuscript editing

LL: performed experiments

OL: performed experiments

LFP: participated in experiment design, assisted with manuscript editing

JCF: assisted with manuscript editing

TSJ: provided key reagents, conducted data analysis

FHF: provided key reagents, conducted data analysis, assisted with manuscript editing

MHH: designed experiments, provided funding, wrote manuscript

